# Dynamic regulation of neuronal vault trafficking and RNA cargo by the noncoding RNA, Vaultrc5

**DOI:** 10.1101/2025.05.16.654162

**Authors:** Mason R. B. Musgrove, Laura J. Leighton, Margaux Lebouc, Wei-Siang Liau, Alexander D. Walsh, Paul R. Marshall, Stephanie M. Heyworth, Adekunle T. Bademosi, Nathalie Hertrich, Qiongyi Zhao, Sachithrani U. Madugalle, Ambika Periyakaruppiah, Xiang Li, Joshua W. A. Davies, Haobin Ren, Hao Gong, Esmi L. Zajaczkowski, Nathalie Dehorter, Frederic Meunier, Marina Mikhaylova, Timothy W. Bredy

## Abstract

Vaults are large ribonucleoprotein complexes of unknown function. Here, we report that the vault-associated noncoding RNA, Vaultrc5, is highly enriched at the synapse and is required for activity-dependent vault trafficking in primary cortical neurons. We have discovered that vaults are comprised of unique populations of coding and non-coding RNA, and that this cargo varies dynamically between subcellular compartments. In addition, Vaultrc5 knockdown at the synapse shifts the RNA cargo toward transcripts associated with immune surveillance, and Vaultrc5 knockdown, *in vivo,* leads to altered fear extinction learning. These findings suggest that the Vaultrc5 is critically involved in coordinating the experience-dependent trafficking of vaults and related RNA cargo, which may represent a novel feature of neuronal plasticity associated with learning and memory.

## 1.0 Introduction

It is well established that neuronal RNAs are differentially expressed and localised in response to activity, particularly those that are active at the synapse (Hirokawa, 2006; Sossin & DesGroseillers, 2006). RNA localisation is required for synaptic plasticity, as it allows for the translation of synaptic RNA pools in response to learning induced synaptic activity that support the formation of memory (Kang & Schuman, 1996; Hirokawa, 2006; Doyle & Kiebler, 2011). However, the selective packaging, synaptic targeting, and translation of synapse-destined RNA is poorly understood.

RNA is typically transported in ribonucleoprotein (RNP) particles called trafficking granules (Kiebler & DesGroseillers, 2000; Steiner et al., 2006). Due to the size of these granules, their formation most likely occurs in the cytoplasm via polyribosomes. Since trafficking granules bind motor proteins on microtubules to move their cargo throughout the cell, how and where trafficking granules form is vital for regulating synaptic plasticity (Antar et al., 2005; Chen et al., 2006; Knowles et al., 1996). Revealing the complete repertoire of synapse-enriched RNAs, how they are trafficked, and when this occurs is therefore paramount to understanding brain function.

Vaults are large 13 MDa ribonucleoprotein complexes that are highly conserved across Eukarya, resemble cages, and transit between the neuron soma and synapse. Major vault protein (MVP) is the main constituent and is vital for vault structure, with protein structure studies implicating pH as an opening signal thanks to MVP’s EF-hand motif (Paspalas et al., 2009; Poderycki et al., 2006; van Zon et al., 2002). Knockout studies have demonstrated that vaults self-assemble completely from MVP, with 78 copies necessary to form its flexible, barrel-like structure (Lodwick et al., 2025; Yu et al., 2017), and MVP knock-out prevents vault formation (Chung & Eng, 2005).

Although the remaining components-TEP1, VPARP, and the small noncoding RNA Vaultrc5, are dispensable for vault structure (Anderson et al., 2007; Kickhoefer et al., 2001; Liu et al., 2004) they are likely critical for the functional activity of vaults. For example, TEP1 is required for Vaultrc5 to associate with the vault (Liu et al., 2004), and VPARP is an uncharacterised poly (ADP)-ribosyltransferase (Poderycki et al., 2005). This suggests that vaults could locally regulate nucleic acids by poly(ADP)ribosylation-consequentially affecting multiple core cellular signalling pathways (Rack, 2021 #1289).

Vaultrc5 is highly structured and the only known RNA component of the vault (Kickhoefer et al., 2003). Previous studies suggest that MVP and Vaultrc5 may be involved in synaptic function. MVP directly associates with AMPA receptors, dendritic mRNAs and ribosomes, and its absence leads to impaired plasticity {Paspalas, 2009 #127}{Ip, 2018 #161}. In addition, both MVP and Vaultrc5 co-immunoprecipitate with proteins involved in the MAPK pathway (Wakatsuki et al., 2021). However, whether these characteristics are directly associated with the function of intact vaults in neurons is not known.

Here, using a targeted RNA degradation platform to knock down Vaultrc5 combined with super resolution microscopy and RNA sequencing, we analysed the functional role of Vaultrc5 in the regulation of vault trafficking and contents in primary cortical neurons, *in vitro*. We find that vaults and Vaultrc5 RNA colocalise at the synapse, and that Vaultrc5 is required for vault trafficking and mobility. Vault immunoprecipitation in activated primary cortical neurons followed by RNA sequencing revealed the unique cargo being trafficked between the nucleus and synapse and Vaultrc5 knockdown dramatically altered the RNA contents of vaults in the synaptic compartment. In addition, we found that Vaultrc5 is highly expressed in the synaptic compartment in the adult brain, and that Vaultrc5 knock-down impairs fear extinction learning. Together, these data indicate that neuronal vaults are RNA trafficking granules that are heavily dependent on the activity of the noncoding RNA Vaultrc5, and suggest a novel mechanism of RNA localisation that may influence plasticity, learning and memory.

## 2.0 Methods

### 2.1 Reagents table

### 2.2 Animals

Wild-type 10-week old C57BL/6J male mice (Australian Research Council (ARC)) were pair-housed in cages with a Plexiglas divider, and maintained on a 12-hour light/dark schedule and given *ad libitum* access to food and water. Testing was conducted during light phase in red light-illuminated testing rooms. All animal use and training, including embryos used for dissection, was in line with the Animal Ethics Committee of the University of Queensland and the Australian Code of Practice for the Care and Use of Animals for Scientific Purposes (eighth edition, 2013).

### 2.3 Construct generation

All constructs used the pFSyn(1.1)GW lentiviral expression vector (Addgene, #27232), which incorporates the rat synapsin 1 promoter, as a backbone. Each construct was verified with Sanger sequencing using custom primers (Supplementary Table S1).

### 2.4 CRISPR-Inspired RNA Targeting System (CIRTS)

The CIRTS cassette was sourced from the Dickinson laboratory at the University of Chicago, U.S.A. This cassette was PCR amplified and inserted between the AgeI and XbaI sites, with an NheI site generated upstream of XbaI (all New England Biolabs). Guide RNAs targeting Vaultrc5 or a scramble control of the respective guide were synthesised at Integrated DNA Technologies, then inserted into the XbaI site (Supplementary Table S1). The constructs were assayed for knockdown efficacy and consistency in primary neurons using a combination of quantitative real-time PCR, and digital PCR.

### 2.5 RT-qPCR & dPCR

One microgram of RNA was synthesised into cDNA using the QuantiTect Reverse Transcription Kit (Qiagen) per the manufacturer’s protocol. qPCR was performed on a RotoGeneQ (Qiagen) real-time PCR cycler with SensiFAST SYBR master mix (Bioline) and appropriate primers. Phosphoglycerate kinase I was used as an internal control. The threshold cycle for each experiment was chosen from the samples’ linear range, and all samples normalised to the internal control using the ΔΔCT method. Each reaction was run in duplicate. Samples were then transferred to the QIAcuity One Digital PCR cycler for absolute quantification in 24-well, 26,000-partition plates (both Qiagen). PCR cycling parameters were identical to qPCR, and analysed as described below.

### 2.7 Vault tagging

Major Vault Protein (MVP) was acquired already-fused at the C-terminus to Green Fluorescent Protein (eGFP) from Origene (MG210985), then inserted into the BamHI site on the pFSyn(1.1) expression vector (Appendix).

### 2.8 Lentiviral production

All constructs were incorporated into individual lentiviral vectors using a third-generation packaging system: pMD2.3 (Addgene #12259), pRSV-REV (Addgene #12253), and pMDLg/pRRE (Addgene #12251). Plasmids were transfected per manufacturer’s protocols using Lipofectamine 3000 (Thermo Fisher Scientific) into HEK293T cells at 80% confluency in culture medium: DMEM (Thermo Fisher Scientific #11965092), 1% GlutaMAX (#35050061), and 1% Penicillin-Streptomycin (#15140122) (both Gibco). After four hours, culture medium was supplemented with 10mM sodium pyruvate (Sigma-Aldrich #P2256), and 10mM sodium butyrate (Sigma-Aldrich #B5887) was added to inhibit cell growth. After fourty-eight hours incubation at 37°C and 5% CO2, virus was collected in 20% sucrose (Sigma-Aldrich #S0389) via ultracentrifugation. Titre was determined using the Lenti-X qRT-PCR Titration Kit (Takara Bio).

### 2.9 Lentiviral delivery & behavioural training Cannulation surgery

Double cannulae (Plastics One) were implanted into the infralimbic prefrontal cortex in the anteroposterior plane along the midline. Implantation locations were centred at +1.8mm in the anteroposterior, and -2.8mm in the dorsoventral plane. Mice were then placed into single housing conditions and given one week to recover prior to behavioural training.

#### Lentiviral infusion

Mice received two lentivirus injections (1μl, 1E7 – 1E8 IU/ml) over 48h. Sterile gas-tight 1700 series syringes (Harvard Apparatus) were used with the Fusion 100 Infusion pump (Chemyx) for this purpose.

#### Behavioural training

Cued fear conditioning and extinction were assayed using two conditioning chambers (Coulbourn Instruments), with two transparent and two stainless steel walls, and a steel grid floor (3.2mm diameter, 8mm x 8mm squares). Contexts A and B were differentiated by spraying context A’s grid with a dilute lemon odour, and covering B’s grid with an opaque plastic surface sprayed with dilute vinegar odour. This was done to minimise context generalisation.

Cameras within the boxes captured movement, which was analysed with FreezeFrame 4. This activity was scored by calculating what percentage of the trial the mouse spent >1s immobile. These scores were transferred to excel and assigned to ‘time windows’ according to which trial was performed- e.g. a pre-conditioned stimulus (pre-CS) timepoint for 120s, followed by the first conditioned stimulus (CS, CS1) for 120s.

The training protocol proceeded as follows, with a 120s pre-CS timepoint before each. For fear conditioning, mice were exposoed to three pairings of a 120s, 80dB, 16kHz tone (CS), which coterminated with a 1s, 0.7mA foot-shock as an unconditioned stimulus (US) in Context A. CS was split by 120s intertrial intervals. The mice were then matched into treatment groups based on freezing scores during CS3. Fear extinction proceeded in context B where mice were initially split into either extinction training (EXT) or retention control (RC) groups. Both groups were habituated for 120s. EXT mice were subject to 60 nonreinforced 120s CS presentations, with 5s intertrial intervals. RC mice were kept in the context for an equivalent time without CS or US. Retention tests followed in context B at 24h and 7d later, with three 120s CS presentations and 120s intertrial intervals.

Behavioural analysis was performed by inferring memory from freezing scores. Animals that did not reach at least 30% freezing by fear conditioning CS3 during fear acquisition were removed, as were those determined to be outliers via analysis in Graphpad Prism 10.

#### Prefrontal cortex extraction

After behavioural training, animals were euthanised via cervical dislocation, and brains immediately extracted and placed on ice. Brains were then sliced into 40µm slices, or dounce-homogenized in a 2ml tissue grinder (Kimble Chase) and processed as appropriate per experiment.

### 2.10 Primary cortical neuron culture

E16 pregnant dams were euthanised via cervical dislocation, embryos extracted and euthanised via immersion in 4°C dissecting medium. Cortical tissue was extracted from embryonic day 16 embryos by removing the skull and meninges with fine-tipped tweezers. Neurons were added to neuronal culture medium: Neurobasal (Thermo Fisher Scientific, #21103049) containing 10% fetal bovine serum (Gibco #26140079) and 1% penicillin-streptomycin, which was supplemented with papain (#P4762) and benzonase (#E1014) (both Sigma-Aldrich). This mixture was incubated at 37°C for twenty minutes before homogenisation via gentle pipetting, then passed through a 40μm cell strainer (BD Falcon). Neurons were plated on well-plates or dishes coated for twenty-four hours with 1.2mM poly-L-ornithine (#P4957), 100mM boric acid (#B0349), 25mM sodium tetrahydroborate (#213462), 75mM sodium chloride (#S9888) (all Sigma-Aldrich) in ultrapure water. Culture medium was added and supplemented with B-27 (Invitrogen #17504), then replaced twenty-four hours later, and neurons were cultured at 37°C and 5% CO2 for seven to fourteen days prior to fixation or RNA extraction. Lentivirus was applied at DIV2 where appropriate.

### 2.11 Synaptosome preparation

Primary cortical neurons at DIV7 from 100mm tissue culture dishes (Falcon) were fixed with 1% methanol-free paraformaldehyde (Thermo Scientific #289206), then dissociated via incubation with StemPro Accutase (Invitrogen #00455556). Neurons were then pelleted in washing buffer, and synaptosomes prepared as previously described(Dunkley et al., 2008). Unlike previous experiments, the sample was split into “cytoplasmic” and “perisynaptic” fractions by pooling and retaining the nuclear and cellular debris fractions generated across the experiment.

### 2.12 Vault immunoprecipitation

Primary cortical neurons transduced with the MVP:GFP, and Vaultrc5 CIRTS lentivirus constructs, were extracted as described above, and either processed as neurons or turned into synaptosomes. Dynabeads Protein G (Invitrogen #10003D) were resuspended in native lysis buffer: 25mM tris-hydrochloride (Roche #10812846001), 150mM potassium chloride (Sigma-Aldrich #P3911), 1% triton X-100 (Sigma-Aldrich #X100), 5mM EDTA (Bioworld #40520000), 5mM DTT (Thermo Scientific #R0861), 2.5ul/ml RNaseOut (Invitrogen #10777019), 1X Halt Protease Inhibitor Cocktail (Thermo Scientific, #78438). Beads were added to the samples and left to rotate for thirty minutes at 4°C. Next, fresh protein G beads were incubated for ten minutes with 5µg of either mouse anti-GFP (abcam #ab6556) or mouse immunoglobin G (Diagenode #C15200001) in native lysis buffer. These bead-bound antibodies were added to the samples and left to rotate for one hour at 4°C. On a magnet, each sample was washed several times with native lysis buffer, then low, and high salt buffers: 0.2X SSPE (Sigma-Aldrich #S2015), 1mM EDTA, 0.05% TWEEN-20 (Sigma-Aldrich #P1379) for low salt, plus 137.5mM sodium chloride for high salt, and 5mM DTT, 2.5ul/ml RNaseOut, 1X Halt Protease Inhibitor Cocktail. After final washes in native lysis buffer, samples were resuspended in 33ul reverse crosslinking buffer: 6% w/v lauroyl sarcosine (Sigma-Aldrich #L5125), 5mM EDTA, 5mM DTT, 1ul/sample RNaseOut, 0.8U/ml proteinase K (New England Biolabs, #P8107), in DPBS (Gibco #14190144). Samples were then incubated for one hour at 42°C, and one hour at 55°C.

### 2.13 RNA extraction

Each sample was incubated with Agencourt RNAClean XP Beads (Beckman Coulter) for ten minutes, allowed to clear on a magnet, then washed twice with 80% ethanol. Samples were dried at 37°C for four minutes, then incubated in ultrapure water for a further ten minutes to retrieve the sample as supernatant. DNase I (Zymo Research, #E1010) was then applied for ten minutes, and samples incubated for ten minutes with fresh RNAClean beads. After clearing on a magnet for ten minutes, a further two washes were performed with 80% ethanol. Beads were then dried and resuspended in ultrapure water as before, and supernatant retrieved. Resulting RNA was quantified using the QuBit 4 Fluorometer (Invitrogen) while the Qubit RNA High Sensitivity Kit, and RNA Integrity Score (RIN) was assessed using the Agilent 2100 Bioanalyser with the Agilent RNA 6000 Pico Kit.

### 2.14 RNA Sequencing

Short-read sequencing was performed in a single lane on an Illumina MiSeq or NovaSeq using the Takara SMARTer Stranded Pico Input Mammalian v2 kit, per the manufacturer’s instructions. Briefly, RNA samples were reverse-transcribed using first-strand cDNA synthesis, ribodepleted, purified using AMPure XP beads (Beckman Coulter), individually barcoded, and amplified for 16 cycles before final purification. Library profiles were analysed on the Agilent Bioanalyser 6000 using the DNA 1000 kit (both Agilent), and then pooled to a single sample at a concentration of 20Nm. An additional purification step was used to remove excess adapter dimer prior to running the sample.

### 2.15 Imaging

#### Immunohistochemistry

Primary cortical neurons plated on coated 35mm glass discs were fixed in 4% w/v paraformaldehyde (Sigma-Aldrich, #158127) for 10 minutes at room temperature, then permeabilised in PBS containing 0.2% Triton X-100 (Sigma-Aldrich, #93443). Samples were washed several times, then blocked using in this buffer plus 10% donkey serum (Sigma-Aldrich, D9663) for one hour at room temperature. Primary antibodies were added at 1:1000, and incubated overnight at 4°C. Samples were washed as before, then incubated with secondary antibodies for one hour at room temperature, and either sealed on microscopy slides using nail polish or stored in PBS at 4°C.

#### Live cell microscopy

Primary cortical neurons plated on coated 35mm dishes were transduced at DIV2 with lentiviruses as appropriate. At DIV7, culture medium was exchanged for buffer containing 15mM HEPES, 145mM sodium chloride, 2.2mM calcium chloride, 0.5mM magnesium chloride, 5.6mM D-glucose (#G8270), 0.5mM ascorbic acid (all Sigma-Aldrich), and 0.1% bovine serum albumin (Invitrogen, #AM2616) in ultrapure water. Neurons were imaged at 37°C and 5% CO2, with and without stimulating with 56mM potassium chloride. Where appropriate, neurons were pre-incubated for ten minutes with YO3-3PEG-Biotin fluorophore (abmgood, #G957).

#### Stimulated emission depletion (STED) microscopy

Neurons plated and fixed on 35mm glass discs as described were imaged dry on a STEDYCON microscope (Aberrior) using an 100X oil objective (1.45 NA). STED utilised pulsed 775nm and 405nm lasers for depletion and excitation, respectively. Detector time gates were set between 0.5-6ns depending on wavelength.

#### Spinning-disc confocal microscopy

Neurons were plated as above were kept in buffer per the live microscopy section at room temperature. These were imaged on an Olympus SpinSR10 Spinning Disk confocal controlled by Zeiss imaging software. Images were taken using the CSU-W1 T2SSR confocal disk unit at 1x zoom with an 100x oil immersion objective (UPLXAPO100X0, 1.45 NA). Z-stacks were generated with a step of 0.5μm, resulting in 15-20 slices each.

#### Superresolution microscopy

Neurons were plated as above and kept as per the live microscopy section were imaged on an Abbelight SAFe 360 inverted single-molecule localisation microscope controlled by Abbelight NEO software. Video was taken using an ORCA-Fusion BT sCMOS cameras with an 100x oil immersion objective (Nikon CFI Apo TIRF, 1.49 NA). Parameters were 16,000 frames in a 256 x 256 format, at approx. 50fps using a 120mW 488nm laser at 30% power.

### 2.16 Experimental design and statistical analysis

STED colocalisation analysis was performed in FIJI as follows. Synaptophysin was used to define an uninterrupted neurite (1 per image, n = 4) for analysis. The Otsu thresholding algorithm for minimising internal variation was used to process each channel along this neurite (e.g. Bassoon, and MVP separately), and converted to a ‘mask’. This mask was used as a reference to determine the total staining in each channel. The masks from both channels were combined to create an ‘overlap mask’, which was measured against each channel to determine the amount of overlapping staining. Overlapping staining was divided by total staining to generate a percentage overlap, then the percentage of MVP overlapping with either marker was analysed by two-way ANOVA.

For all short-read RNA sequencing experiments, data was processed as follows: Cutadapt (v3.7) was used to clip low-quality nucleotides (Phred < 20) and adapter sequences. Next, HISAT2 (v2.2.1) was used to align the reads to the mouse genome (mm10), with HTSeq-count (v2.0.5) to generate the count matrix for each sample. Finally, edgeR (v4.4.1) was used to analyse differential expression within groups.

The fear learning experiment was carried out with twelve animals per group, as power calculations within our previous experiments demonstrated that a minimum of eight animals was sufficient to determine statistical significance. Twelve mice therefore account for removal due to outliers, surgical complications, or inadequate fear acquisition. This experiment was repeated, and animals pooled across both instances to avoid a single cohort effect.

Previous sequencing experiments have detected effect sizes of 1.2-1.5 fold with n = 6 samples per treatment. As we were unable to reach this in all sequencing experiments, this is discussed in this limitations section. For single molecule tracking, kymographs were generated with an n = 10 per group (one per dish) to minimise within-group variation. Each kymograph was constructed from approximately 12µm of dendrite, at a thickness of 1µm. Statistical analyses were generally one- and two-way ANOVAs accompanied by a post-hoc comparison between all groups and the relevant control group. Where two-way ANOVAs detected a significant main interaction effect, one-way ANOVA followed by Šidák’s post-hoc analysis were performed.

## 3.0 Results

### 3.1 Vaults are abundant in both the pre-and post-synaptic compartment, and their trafficking depends on Vaultrc5

Using STED microscopy to analyse the trafficking of Vaults in primary cortical neurons, in vitro, we measured the distribution of major vault protein MVP, and examined its colocalization with presynaptic and postsynaptic markers. Pre-synaptic terminals were identified using Bassoon (Fig 1A-D), and post-synaptic terminals with Shank3 (E-H). For neurons stained with Bassoon (Fig 1A), images were taken using the axon terminal as a guide, while Shank3 (Fig 1E) used dendrites. We found that MVP was biased towards the presynaptic terminal, which, given Vaultrc5’s expression characteristics (Fig 1A, B), could indicate that vaults spend most of their time transiting from the nucleus to the synapse.

**Figure 1.**
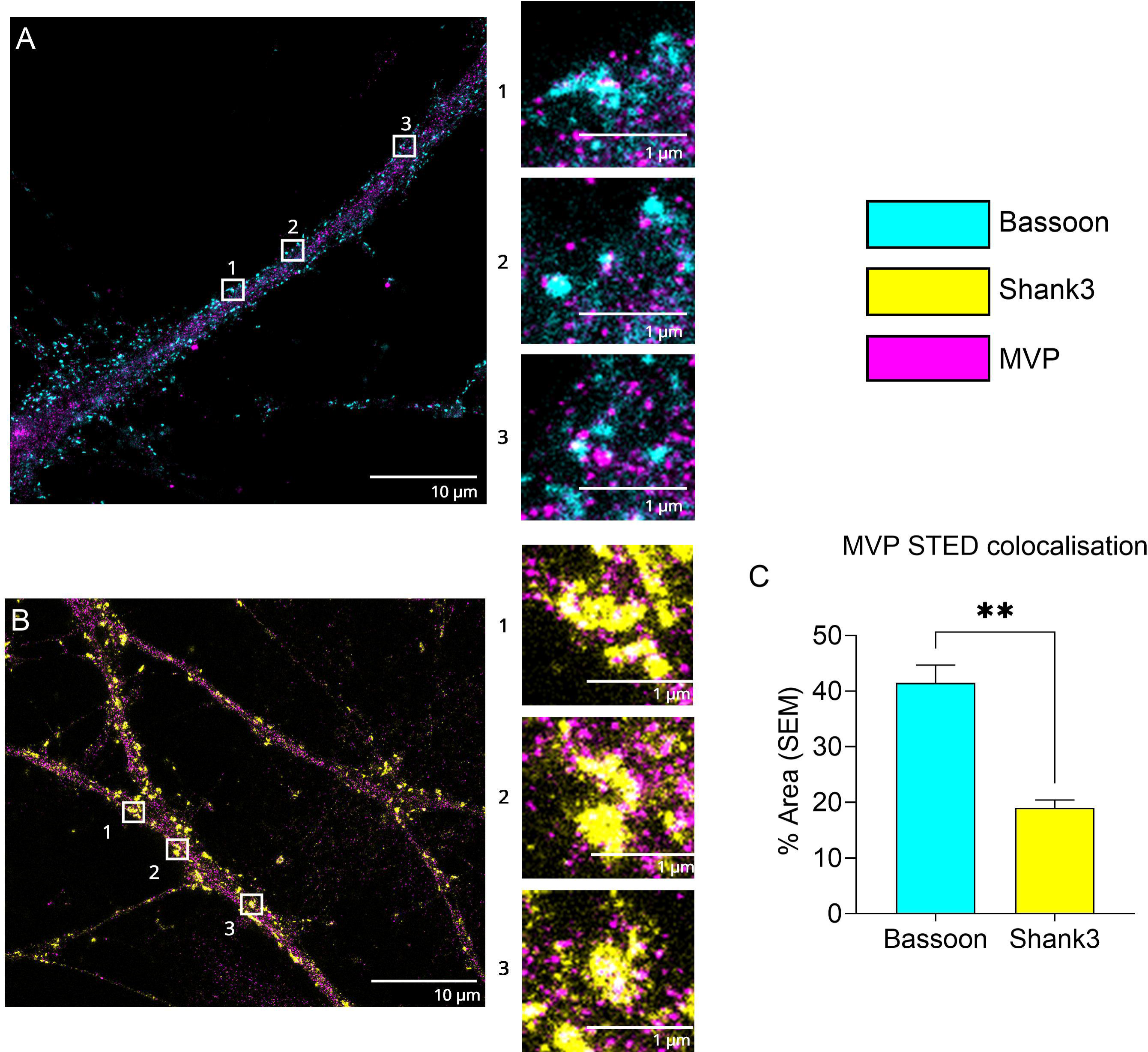

We then used an MVP fused to GFP (MVP-GFP) fusion protein to tag vaults, and tracked them in live cells using kymography with a spinning-disc confocal microscope for 3 minutes at 2 frames per second (Fig 2A). Neurons were transfected with either the GFP tag, or the GFP tag plus a knockdown construct targeting Vaultrc5, and imaged under 20mM KCl-induced neuronal depolarisation conditions (Supplementary Figure S1). After imaging, 12µm sections were processed using the KymoClear 2.0 plugin for FIJI, splitting molecular movement into anterograde, retrograde, and pausing behaviour (Fig 2B). Vaultrc5 knockdown led to a significant reduction in events per 10µm (Fig 2C). Knockdown did not impact run time per movement (Fig 3B), while pausing remained the most common, indicating that the effect on events per 10µm was not due to shorter movements. Velocity, and colocalization with pre- and post-synaptic markers was not affected (Supplementary Figure S2).

**Figure 2.**
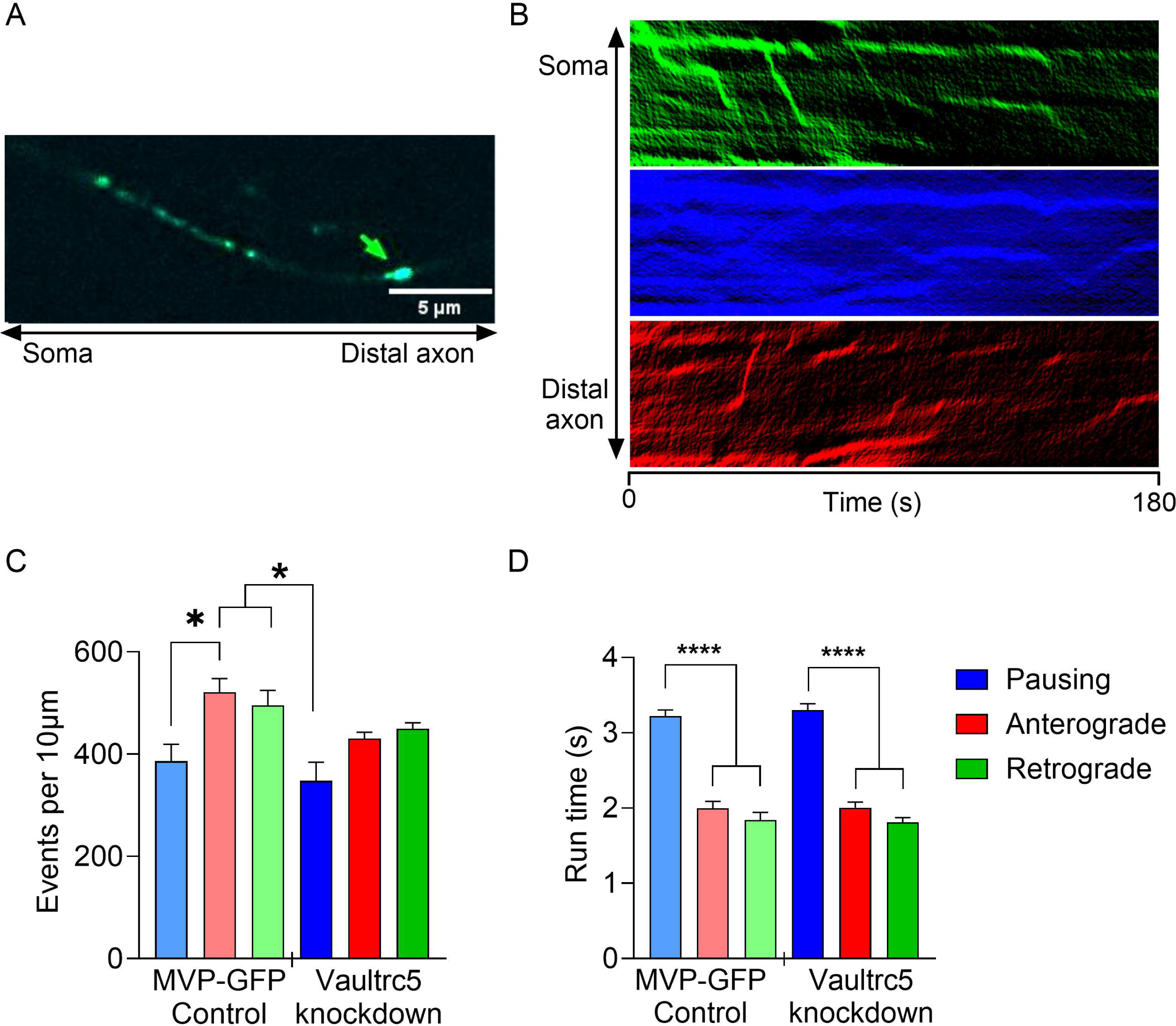

**Figure 3.**
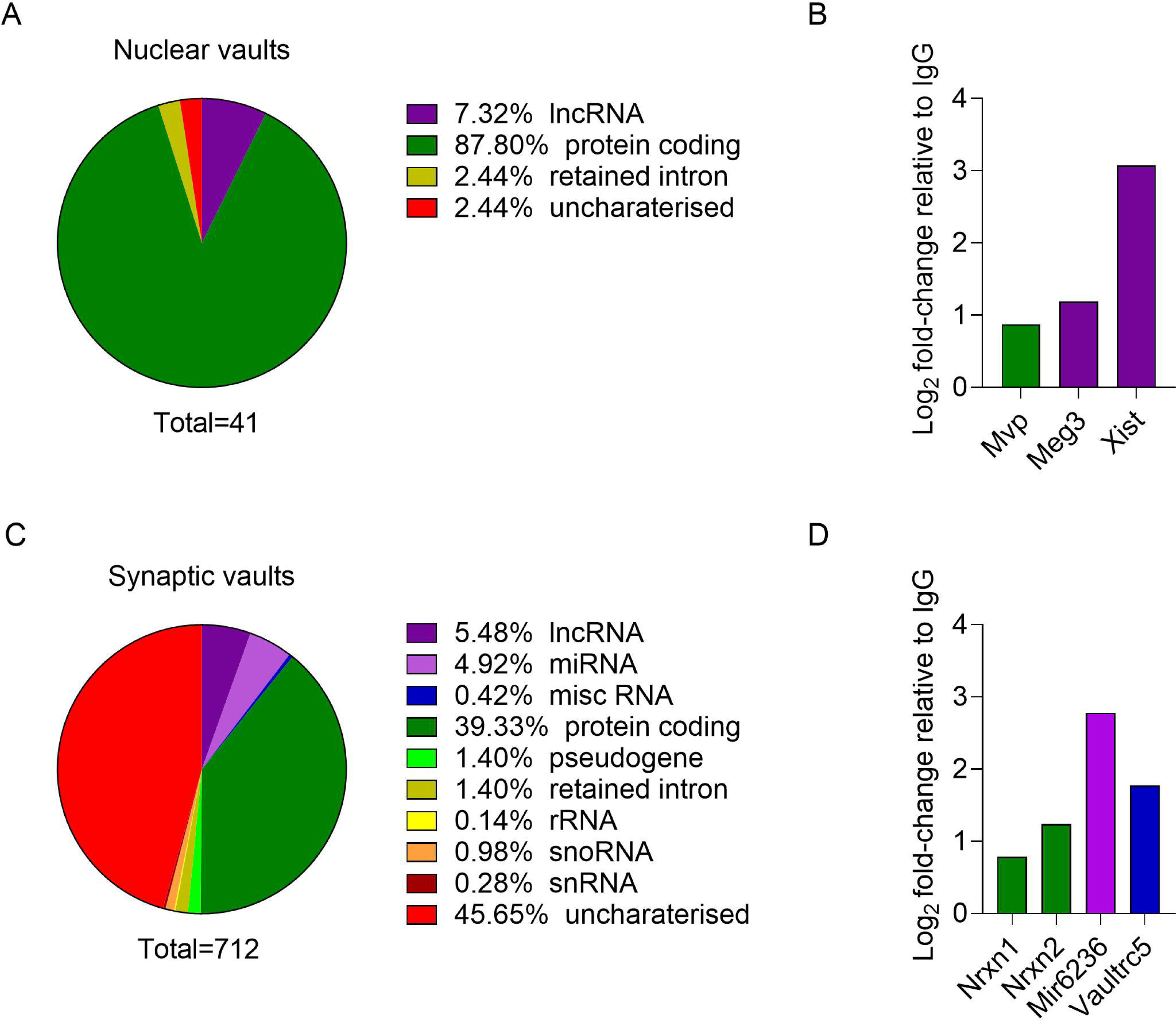

### 3.2 Vaults are trafficking granules

Having confirmed that vaults are highly mobile in neuronal processes and that trafficking relies on Vaultrc5, we then examined the effect of Vaultrc5 on the vault cargo in the nuclear and synaptic compartment. To address this, we immunoprecipitated vaults from nuclear and synaptosome fractions derived from KCl-activated primary cortical neurons and performed RNA sequencing on the vault-enriched fractions (Supplementary Figure S3

Nuclear vaults (Fig. 3A, C) contained primarily protein-coding transcripts, with small populations of lncRNAs, and unknown or uncharacterised transcripts. Most transcripts were involved in intercellular signalling and the immune response, including Meg3, a key lncRNA involved in learning, memory, and the immune response (Yi et al., 2019); Xist, required for X-chromosome dosage compensation(Sahakyan et al., 2018); and the major vault protein mRNA.

Synaptic vaults (Fig. 3B, D) had a more than ten-fold increase in transcript diversity, with cargo split between uncharacterised and protein coding, with the majority being unknown. An array of smaller transcript populations was also present, primarily lncRNA and miRNA. Known transcripts included several involved in synaptic plasticity, such as Nrxn1 and 2, and miR6236. The neurexin family are well-known presynaptic cell adhesion proteins present throughout the nervous system (Lin et al., 2023), while miR6236 was recently shown to regulate neuronal repair and development (Chen et al., 2021). This is consistent with the idea that vaults may play a role in synaptic modulation by trafficking transcripts required for the synapse to dynamically respond to environmental stimuli.

### 3.3 Vaultrc5 regulates the content of synapse-enriched vaults

Considering the increase in transcript diversity within vaults enriched within the synaptic fraction, we examined the effect of Vaultrc5 knockdown on vault cargo at the synapse. Vaultrc5 knockdown dramatically altered transcript diversity, with unknown, and immune signalling-related transcripts becoming more abundant. In addition, Vaultrc5 knockdown led to the accumulation of many unaligned transcripts, the most abundant of which partially corresponded to ribosomal, and mitochondrial RNA (Supplementary Figure S4).

Most transcripts present in synaptic vaults (Figure 4A, C) were mRNAs involved in cellular metabolism. Amongst these were Mtn2, neuroprotective in stroke (Sozmen et al., 2019); Fzd4, involved in dendrite morphogenesis (Bian et al., 2015); and Crlf1, a neurotrophic cytokine receptor ligand (Looyenga et al., 2013). We also detected miR-540, which targets mRNA encoding voltage-dependent calcium channel, and is involved in stress-induced depression (Ma et al., 2016) and BC1, a lncRNA that regulates neuronal mRNA and that appears to be important for spatial learning (Chung et al., 2017). Interestingly, Vaultrc5 was only detectable in the synaptic fraction (Figure 4D).

**Figure 4.**
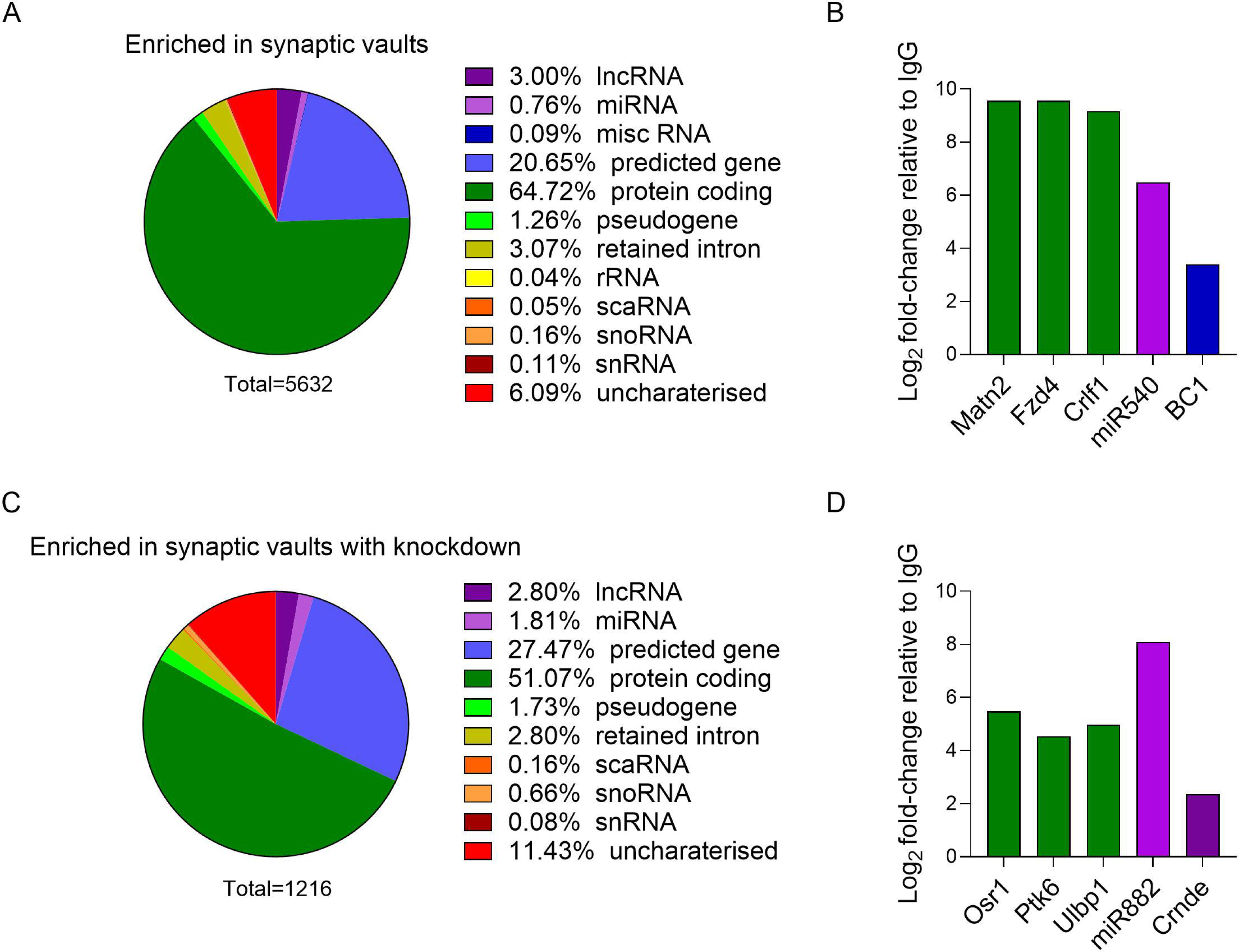

Following Vaultrc5 knockdown (Figure 4B, D), transcripts in synaptic vaults shifted toward unknown and uncharacterised examples, with the most prevalent being associated with immune signalling, including Osr1, a key transcription factor in nervous system development (Nugent et al., 2012); Ulbp1, a positive regulator of the adaptive immune response (Ruan et al., 2022); and Ptk6, a receptor tyrosine kinase involved in the innate immune response (Vlajic et al., 2024). With respect to ncRNA, there was miR-882, a promiscuous miRNA with many cell-cycle and cancer-related targets- one of which is Rev-erbα, a core component of the circadian clock (Griffin et al., 2019; Tian et al., 2021); and Crnde, which regulates the mTOR complex in cell growth and proliferation (Wang et al., 2015). Vaultrc5 was no longer detectable following VaultRC5 knockdown.

The variation in transcript diversity suggests that Vaultrc5 knockdown alters the synaptic RNA pool by biasing it toward processes involved in environmental signalling and the immune system, possibly by influencing the subcellular compartmental distribution of vaults, or by affecting the transcripts being loaded within the Vaults. This would agree with precious work showing that MVP is upregulated in response to environmental signaling across infection, inflammation, and cancer (Mossink et al., 2003; Steiner et al., 2006)

### 3.4 Vaultrc5 is highly enriched at the synapse, *in vivo*

A re-analysis of our previously published *in vivo*, synapse-enriched, RNA-seq data (Liau et al, 2023) revealed a significant population of ncRNAs in the adult prefrontal cortex that are expressed in a subcellular compartment-specific manner (Figure 5A, B). Amongst the ncRNAs localised to the synapse, Gas5, Malat1, Rny1 and Rny3, and Vaultrc5 were the most abundant. We have previously demonstrated that synapse-enriched Gas5 and Malat1 are integral for fear extinction memory (Liau et al., 2023; Madugalle et al., 2023). Rny1 and Rny3 have not been implicated in synaptic function, but have recently been shown to be glycosylated, and present on the outer cellular membrane {Flynn, 2021 #425}.

**Figure 5.**
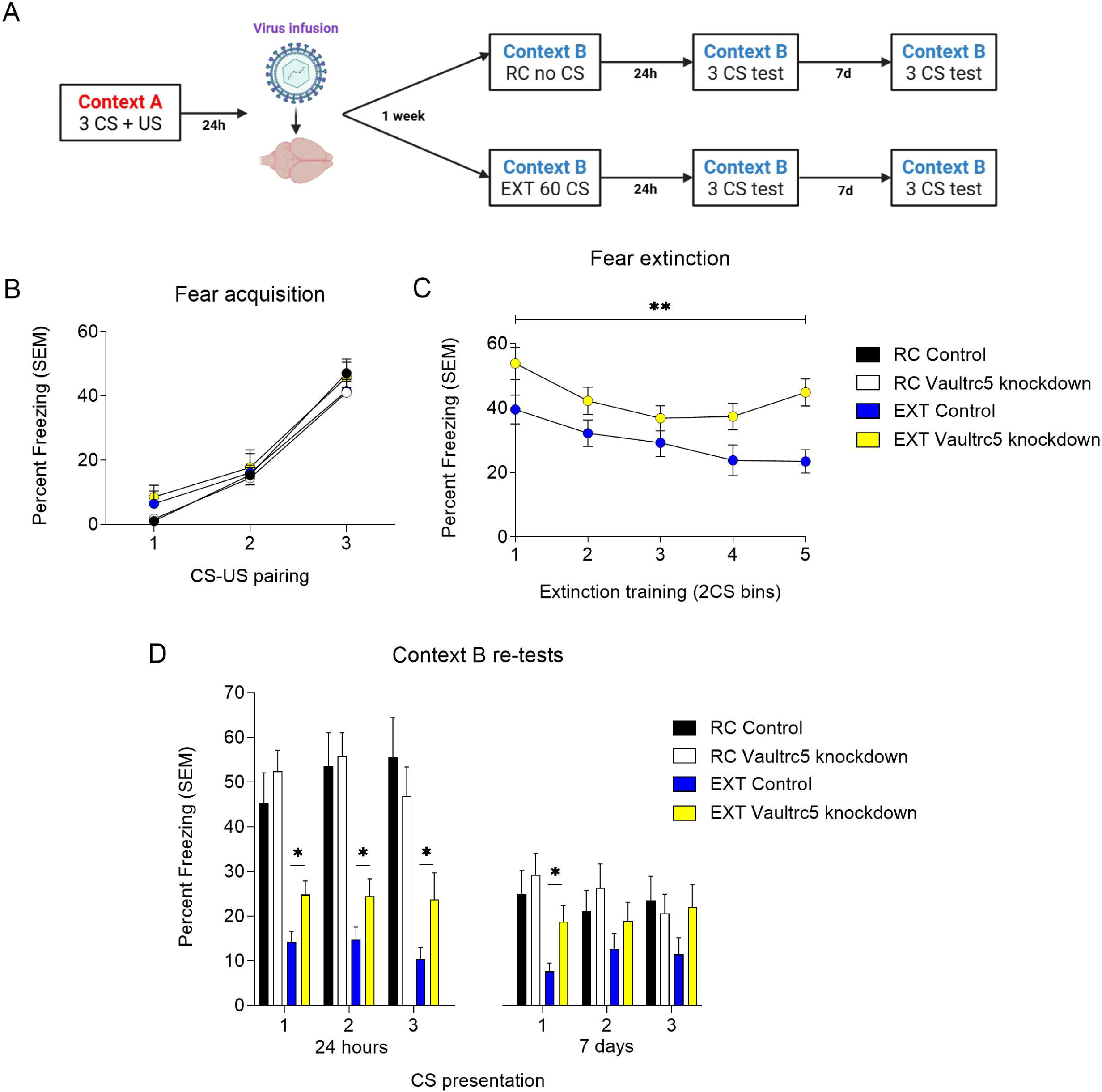

Supporting the vault sequencing data, Vaultrc5 expression was amongst the most dynamically expressed of all synapse-enriched ncRNAs. In retention control mice it was eight times higher in the synapse than in the nucleus-a difference which was not present during fear extinction (Figure 5A, C). These data suggest a role for Vaultrc5, and the accompanying ‘vault’ ribonucleoprotein complex, in the regulation of synaptic plasticity and memory.

### 3.5 Targeted Vaultrc5 knockdown impairs fear extinction learning

To determine whether Vaultrc5 knockdown influences fear extinction, we used a lentiviral vector to deliver the Vaultrc5 knockdown construct directly to the infralimbic prefrontal cortex (ILPFC) 24hr after fear conditioning (Figure 6A). We chose to examine fear extinction because we have previously demonstrated that RNA-mediated mechanisms in the ILPFC are critically involved in this important form of fear-related learning{Liau, 2023 #1290}{Madugalle, 2022 #1139}. All mice acquired a robust response to fear conditioning (Figure 6B). One week after viral transfection, the mice underwent either a 60CS fear extinction (EXT) training protocol in a novel context (Cxt B) or were exposed to Cxt B without CS exposure for an equivalent period (retention control, RC). A two-way repeated measures ANOVA revealed a significant interaction between group and time, with Vaultrc5 knockdown mice exhibiting higher levels of freezing across the first 10 CS (Figure 6C). 24h later, mice treated with Vaultrc5 knockdown showed impaired extinction memory across all three CS exposures (Figure 6D, left panel) and this effect persisted for 7 days (Figure 6D, right panel). There was no effect of Vaultrc5 knockdown on retention control mice indicating that the effect was not due to a disruption of the original fear memory.

**Figure 6.**
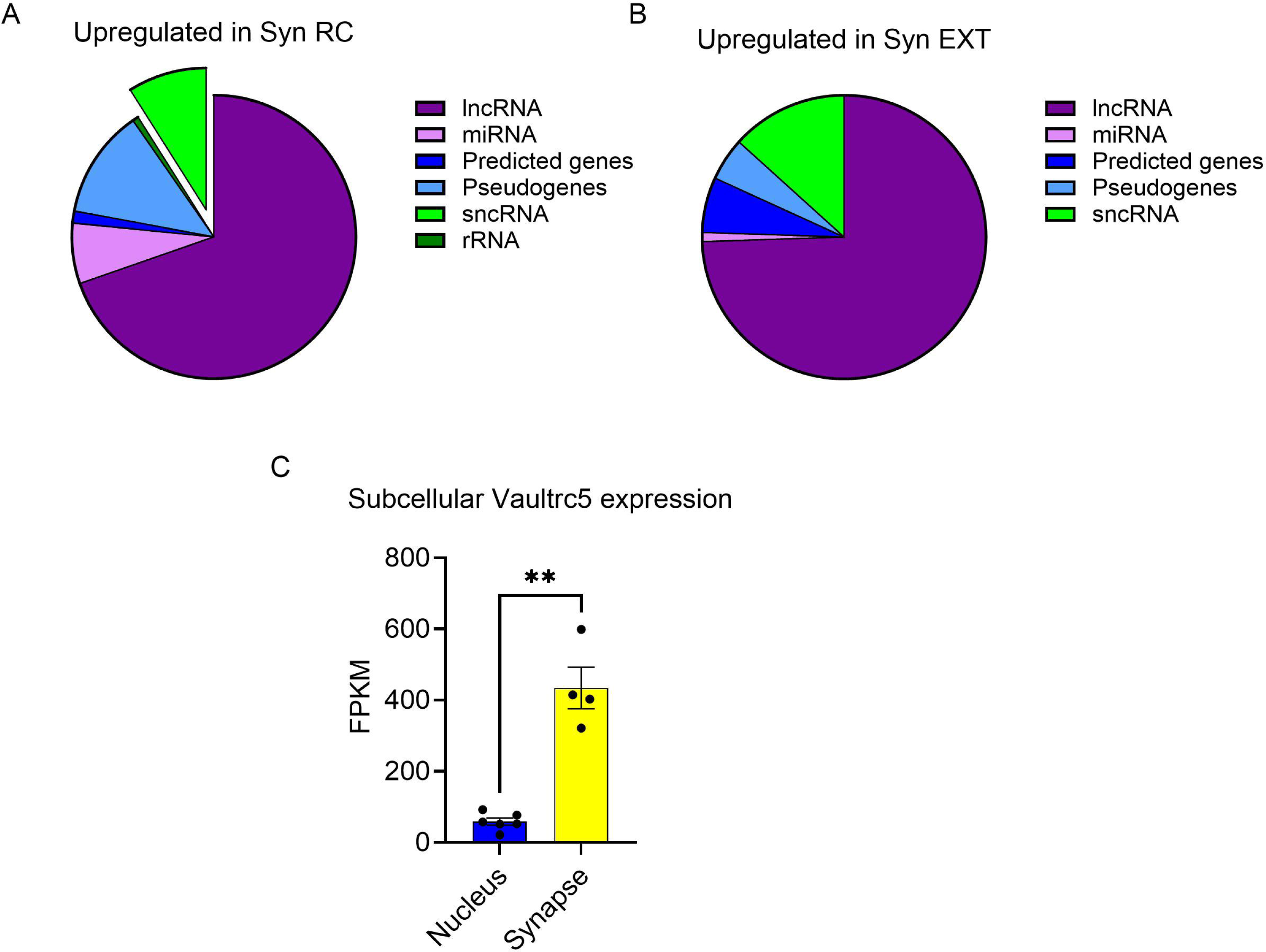

## 4.0 Discussion

Since their discovery in 1986, vaults have been considered to be large, empty, nanoparticles that could be useful as drug delivery vehicles (Casanas et al., 2012; Kedersha & Rome, 1986). In this study, we have demonstrated that vaults are far more complex than previously assumed. We have found that 1) vaults are trafficking granules that shuttle RNA between the soma and synapse, a process that requires the noncoding RNA Vaultrc5; 2) Knocking down Vaultrc5 causes the vault’s cargo to switch to a population of more metabolic, and immune signal-responsive transcripts; and 3) Vaultrc5 knockdown impairs fear-related learning. Taken together, these data indicate that neuronal vaults and Vaultrc5 respond dynamically to environmental cues, and are integral to the regulation of RNA metabolism underlying learning and memory.

The finding that vaults are trafficking granules in neurons is the first evidence of vault function in any context. Early studies on vaults had not examined their RNA cargo due to technological limitations, so they were assumed contain only a few proteins. Further, previous studies had linked MVP and Vaultrc5 to the immune system and synaptic plasticity, but were unable to connect these characteristics to vault function. Here, we show that vaults are present in both the pre- and post-synaptic compartment, albeit with a bias toward presynaptic trafficking, and that the vault transcript cargo changes in response to its environmental context, including the subcellular compartment.

Synaptic compared to nuclear vaults had a far more diverse transcriptome, with the majority being uncharacterised RNAs. Nuclear vaults primarily contained mRNA, and several kinds of ncRNA. These included the MVP mRNA, Meg3, and Xist, all of which are extensively implicated in intercellular signalling and the immune system. Some of the most common transcripts in synaptic vaults were miR-6236, and Neurexin 1 and 2. MiR-6236 was recently shown to regulate neuronal repair and development, while the neurexin family are presynaptic adhesion proteins well-characterised as key to synaptic function. The shift in content and function, and the presence of these specific transcripts, confirm that vaults function as trafficking granules in synaptic modulation.

Our data indicating that Vaultrc5 is required for both vault movement and cargo ties into previous studies indicating that Vaultrc5 has at least two synaptic functions: 1) It is associated with the SQSTM1 complex in autophagy, and 2) both Vaultrc5 and MVP are critical for MAPK/ERK activation at the neurite in synaptogenesis. Here we show that Vaultrc5 knockdown in synaptic vaults leads to a marked accumulation of unaligned reads, unknown transcripts and RNA involved in metabolic, and immune-related signalling, including Ulbp1, Ptk6, and miR-882. Ulbp1 and Ptk6 are regulators of the adaptive, and innate immune response, respectively. miR-882 is a promiscuous miRNA with many cell cycle-related targets-one of which was Rev-erbα, key to the circadian clock and a regulator of inflammation. The change in synaptic vault cargo towards immune signalling-related RNAs suggests that Vaultrc5 acts as a “gear shift”, with its downregulation perhaps causing vaults to become more receptive to environmental signals.

This would fit with the altered movement and extended pausing behaviour of vaults caused by Vaultrc5 knockdown. Thus, vaults appear to monitor, and respond to environmental changes by substituting the cargo they deliver to the synapse. This surveillance function of the vault could be mediated by Vaultrc5 itself, or other miRNAs. miRNA are well-characterised as immune rheostats-fine-tuners of the cellular environment, where small changes in their expression, or binding partner accessibility, has large and varied downstream, and recursive upstream effects. One prominent example is miR-181, which is also within synaptic Vaults, and controls early natural killer T cell development (Henao-Mejia et al., 2013). Interestingly, Vaultrc5 undergoes NSUN2 methylation-dependent, DICER-mediated processing into several miRNA-like small Vault RNAs (svtRNAs). These svtRNAs have been shown to act like miRNA in immune function and disease-for example, svtRNA2-1 is upregulated early in Parkinson’s disease, perturbing cellular metabolism (Minones-Moyano et al., 2013). The signal for their production has not been elucidated but, given our data on synaptic vaults following Vaultrc5 knockdown, their presence could be due to Vaultrc5 fine-tuning the vault to respond to environmental signalling.

The interaction between the vault and Vaultrc5 might also contribute to behavioural phenotypes. Vaultrc5 knockdown prior to fear extinction affected the rate at which mice were able to dissociate the conditioned and unconditioned stimulus, both during within-session extinction training and in the context B tests. Vaultrc5 knockdown likely attenuated the formation of the new, fear extinction trace; however, in order to confirm this, future experiments will examine the effect of Vaultrc5 knockdown prior to fear acquisition, and with knockdown localised to the prelimbic prefrontal cortex.

In the event that these data are due to a disruption of new learning, and given that Vaultrc5 enables the vault to respond to environmental signalling, knockdown may have reduced the intrinsic excitability of neurons prior to fear extinction. This would hinder the brain’s ability to create a new trace. Reduced excitability would also indicate that the vaults’ response of switching to metabolic and immune-related transcripts lead the neurons away from an activated state, focusing instead on processes involved in environmental surveillance.

A reduction in intrinsic excitability could arise from the fact that both Vaultrc5, and MVP interact with the MAPK/ERK pathway at the synapse. First, the EF-hand motif present on MVP is generally known to interact with both pH, and intracellular Ca^2+^, with pH being implicated in Vault opening (Lin et al., 2021; Querol-Audi et al., 2009). Vaultrc5 activating the MAPK pathway could increase MVP’s phosphorylation levels, providing another trigger for Vault opening other than synaptic transmission. Fewer open Vaults combined with less MAPK/ERK activation, leading to decreased synaptic plasticity (Bluthgen et al., 2017), could result in an overbalance of Vaults carrying immune signalling-related transcripts.

In summary, we have discovered that vaults are trafficking granules involved in moving RNA cargo between the neuron soma and synapse, which depends on the activity of Vaultrc5. Vaultrc5 may thus act as a gear-shift to fine-tune the vault’s response to environmental signalling, with potentially important effects on plasticity underlying learning, and memory.

## Acknowledgements

We gratefully acknowledge grant support from the Brain and Behavioral Research Foundation (NARSAD Independent Investigator Award, T.W.B.) and NHMRC Ideas Grant (GNT2003414) to T.W.B. L.J.L. S.U.M. and E.L.Z. were supported by a Westpac Future Scholarship and the University of Queensland. M.M. was supported by the Australian Research Training Stipend. We thank the Queensland Brain Institute Advanced Microscopy Facility for its support and Ms. Rowan Tweedale for manuscript editing.

The authors declare that they have no conflict of interest.

Data are available online at DOI: 10.17632/ckpxd8xx22.1.

---

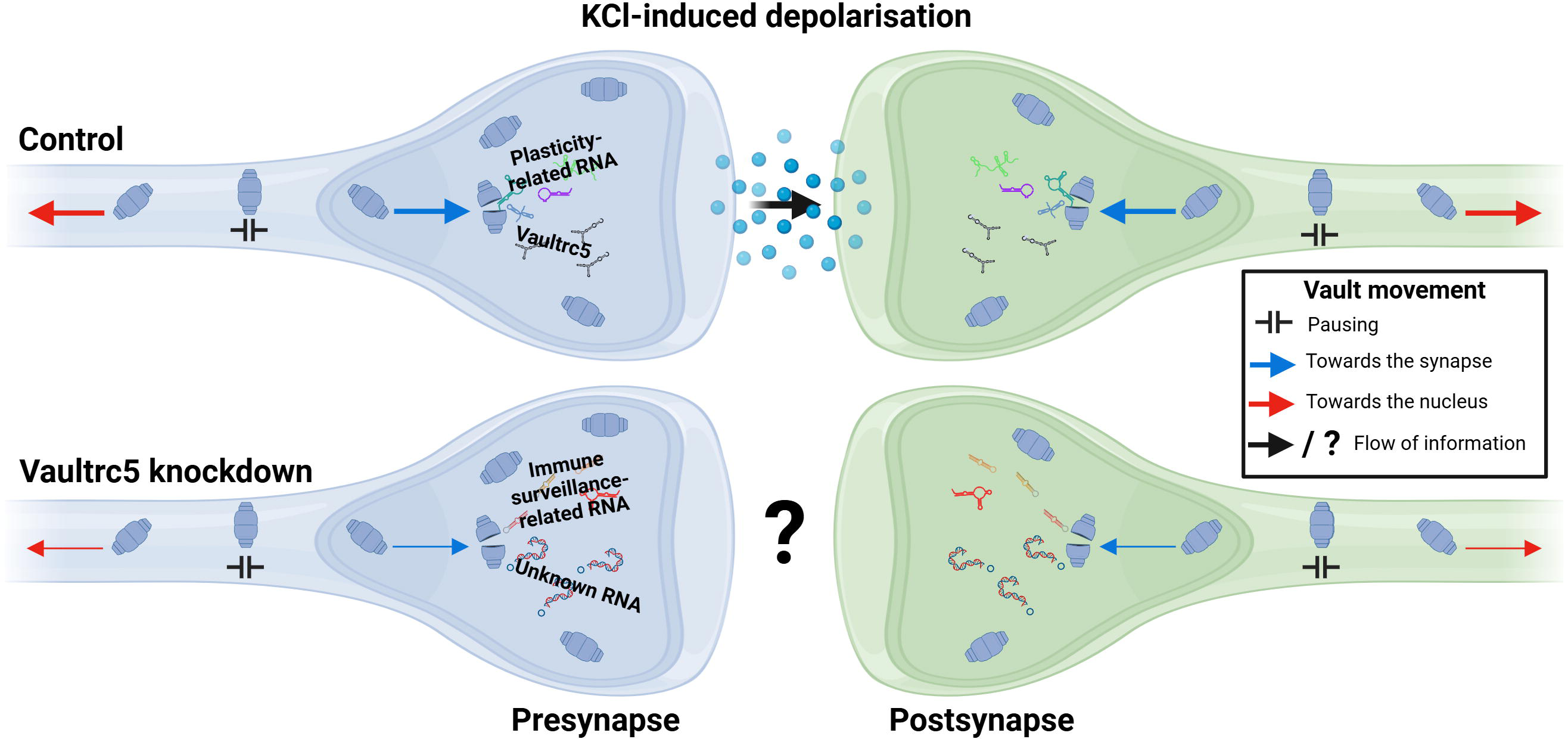

