## Supplementary material for "Dynamic regulation of neuronal vault trafficking and RNA cargo by the noncoding RNA, Vaultrc5": Figure captions

**Figure 1 – MVP is presynaptically enriched.** STED microscopy on DIV14 mouse primary cortical neurons. A) Representative presynaptic STED imagery using Bassoon (teal) and Major Vault Protein (MVP)(magenta). B) Representative postsynaptic STED imagery using Shank3 (teal) and MVP (magenta). C) Percent area of MVP colocalised with either Bassoon or Shank3. Unpaired two-tailed t-test with Welch’s correction. Significantly more MVP was present at the presynapse, with a mean difference of 22.42 ± 4.66% (t = 6.349, df = 4.068, p = 0.0029).

**Figure 2 – Vault trafficking is attenuated by Vaultrc5 knockdown.** Mouse primary cortical neuron (DIV7) kymographs of GFP-labelled vaults, with and without Vaultrc5 knockdown. Particles were tracked using a spinning-disk confocal for 3 minutes at 2 frames per second. One kymograph was generated per cell (n = 10). Videos were analysed using KymoClear 2.0 in ImageJ, then with one- and two-way ANOVA. A) Representative image taken from video used for kymograph in B. B) Representative kymograph of Vault trafficking. Anterograde (red), Pausing (blue), Retrograde (green). C) Events per 10µm per movement vector. Each event was one full path of a tracked molecule or molecular cluster. Pausing behaviour in the control group was significantly rarer than anterograde at 135.3 ± 57.79 events (p = 0.011). Pausing in the knockdown group was significantly less frequent than both control anterograde (164.6 ± 59.89, p = 0.002) and retrograde movement (138.5 ± 59.88, p = 0.013). D) Run time (s) per event. Both groups spent the most time pausing, at an average of 1.301 (p < 0.0001) for control, and 1.395 (p < 0.0001) for Vaultrc5 knockdown.

**Figure 3 – Vault RNA cargo differs between the nucleus and synapse.** Short-read sequencing reads (log_10_CPM) of RNA derived from Vaults per subcellular compartment (n = 4 per compartment) under activated conditions. A, B) Transcripts found in vaults derived from the nuclear compartment (n = 41), and example candidates: Mvp (q = 0.014), Meg3 (q < 0.001), Xist (q < 0.001). C,D) Transcripts found in vaults derived from the synaptic compartment (n = 712), and selected candidates. Nrxn1 (q < 0.001), Nrxn2 (q < 0.001), miR-6236 (q < 0.001), Vaultrc5 (q < 0.001).

**Figure 4 – Vaultrc5 acts as a “gear-shift” for vault function.** Short-read sequencing reads (log_10_CPM) of RNA derived from vaults per subcellular compartment (n = 4 per compartment) under activated conditions. Reads were analysed using pipeline described in methods, and background (IgG) subtracted from Vault immunoprecipitation. A,B) Transcripts enriched in synaptic vaults (n = 5632 transcripts), with example transcripts: Matn2 (q < 0.001), Fzd4 (q < 0.001), Crlf1 (q < 0.001), miR-540 (q < 0.001), Bc1 (q < 0.001). C,D) Transcripts enriched in synaptic vaults following Vaultrc5 knockdown (n = 1216 transcripts), with examples. Osr1 (q = 0.005), Ptk6 (q < 0.001), Ulbp1 (q < 0.001), miR-882 (q < 0.001), Crnde (q = 0.048).

**Figure 5 – Vaultrc5 is subcellularly localised in response to fear extinction learning.** RNAseq data from fear-conditioned (EXT) or retention control (RC) mice, separated according to nuclear or synaptic (Syn) subcellular fraction. A-B) Classes of non-coding RNA found significantly upregulated at the synapse following (A) fear conditioning (retention control, RC) or (B) fear extinction learning (EXT). C) Read count (FPKM) of Vaultrc5 at the nucleus versus synapse. Unpaired two-tailed t-test with Welch’s correction. Vaultrc5 is significantly upregulated in the synapse (t = 6.321, df = 3.173, p = 0.0068).

**Figure 6 – Vaultrc5 knockdown impairs fear extinction learning.** A) Schematic of fear condition and extinction learning paradigm. Mice were treated with either the control (GFP) or Vaultrc5 knockdown construct (n = 16-18 per group across 2 cohorts, 4 groups). B) No significant difference between the groups by the 3rd CS-US of fear acquisition. C) Vaultrc5 knockdown significantly attenuated fear extinction learning across the first 10 CS exposures (F_12,220_ = 2.543, p = 0.0037). D) Vaultrc5 knockdown led to impaired fear extinction memory 24h (2-way repeated measures ANOVA, F_6,104_ = 2.343, p = 0.0366) and 7 d after extinction training (F_6,100_ = 2.817, p = 0.0142). Simple effects analysis revealed a main effect of training on fear extinction memory at CS1 (F_3,52_ = 15.92, p < 0.0001; Šídáks post-hoc analysis, EXT control vs EXT Vaultrc5 knockdown, *p<.05), CS2 (F_3,52_ = 16.41, p < 0.0001, Šídáks post-hoc analysis EXT control vs EXT Vaultrc5 knockdown, * p<.05), and CS3 (F_3,52_ = 10.70, p < 0.0001, Šídáks post-hoc analysis, EXT control vs EXT Vaultrc5 knockdown, *p<.05); and during CS1 (F_3,50_ = 5.549, p = 0.0023, Šídáks post-hoc analysis, EXT control vs EXT Vaultrc5 knockdown, *p<.05) 7 d after training. (Data are presented as percent time spent freezing ± SEM. CS = conditioned stimulus; US = unconditioned stimulus; RC = retention control, EXT = extinction).
