## Supplementary Figure 1 for "Dynamic regulation of neuronal vault trafficking and RNA cargo by the noncoding RNA, Vaultrc5"

**Supplementary Figure X: CIRTS was targeted to Vaultrc5 using thermodynamic structure prediction.** A) CIRTS gRNA probes against structurally predicted conserved regions on the long and short forms of Vaultrc5. Structural predictions were sourced from RNAFold. Probes were designed based on annotation data from the UCSC genome browser. Red bases indicate the highest probability of the predicted structure occurring, while green bases indicate lower probability. B) Data are the average fold-change and standard error from qPCR on primary neuron cDNA after viral transduction of the CIRTS construct targeting Vaultrc5 or GFP (n = 5). All data were normalised first to the reference gene PGK, and then to the GFP condition. CIRTS transduction (hereafter referred to as Vaultrc5 knockdown) significantly reduced Vaultrc5 expression in vitro by 0.47 ± 0.06 fold (F_4,4_ = 1.057, p < 0.0001).

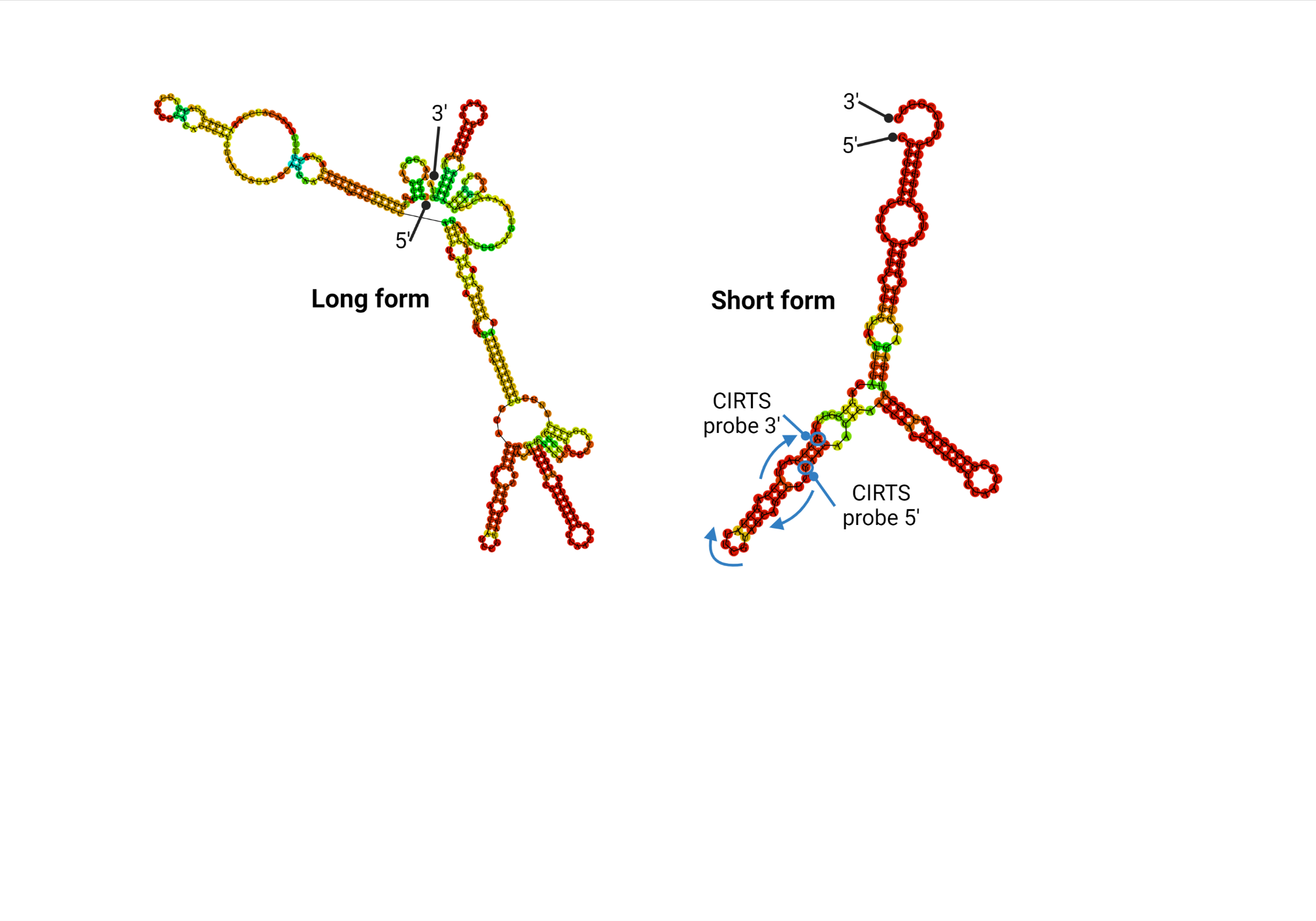

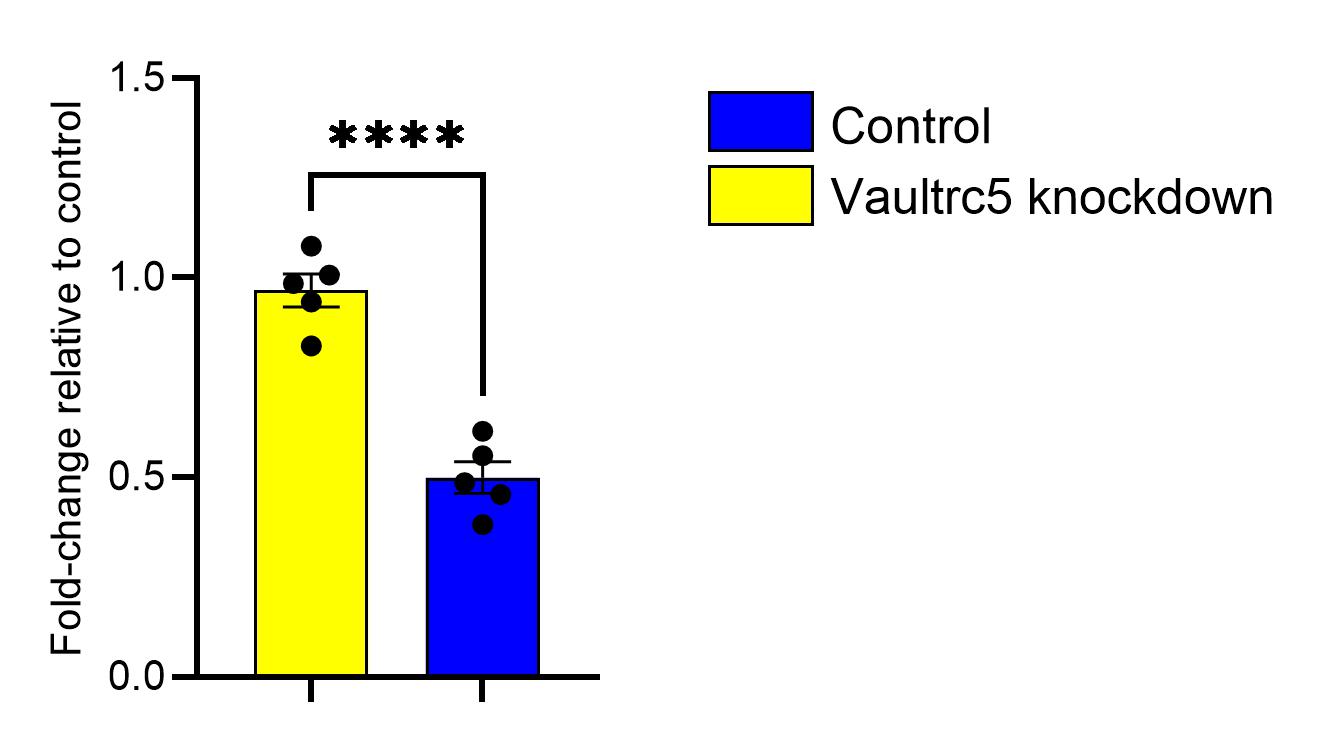
