## Supplementary Figure 2 for "Dynamic regulation of neuronal vault trafficking and RNA cargo by the noncoding RNA, Vaultrc5"

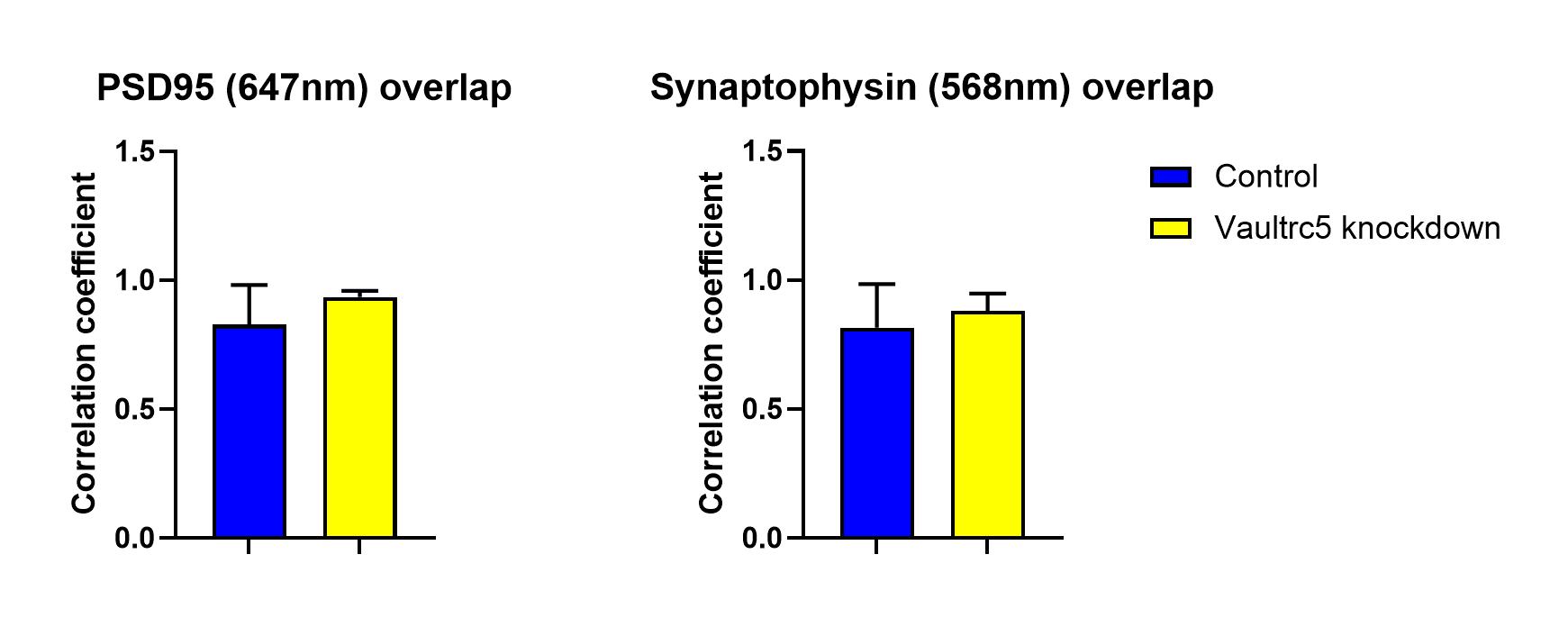

**Supplementary Figure X - Vaultrc5 knockdown didn’t affect colocalization of Vaults with A) PSD-95 or B) synaptophysin.** Data are analyses performed on images taken on a spinning disc confocal, using synaptophysin for presynaptic markers and PSD-95 for postsynaptic. Colocalisation analysis was performed using the BIOP JACOP plugin on FIJI.

**A	B**
