## Supplementary Figure 3 for "Dynamic regulation of neuronal vault trafficking and RNA cargo by the noncoding RNA, Vaultrc5"

**Supplementary Figure X – qPCR on nuclear and synaptic fractions prior to vault immunoprecipitation.** PGK1 was used as a control in both compartments, Polr2a was the nuclear reference gene, PSD95 was the synaptic reference gene (n = 3-4). Analysed via unpaired t-test with Welch’s correction. A) Polr2a was significantly lower in the synaptic compartment (F_3,2_ = 1274, p = 0.0009). B) PSD95 was significantly lower in the nuclear compartment (F_2,3_ = 239.7, p = 0.0241). C) Vaultrc5 was significantly more enriched in the synaptic, than nuclear compartment (F_2,3_ = 53.20, p = 0.0364).

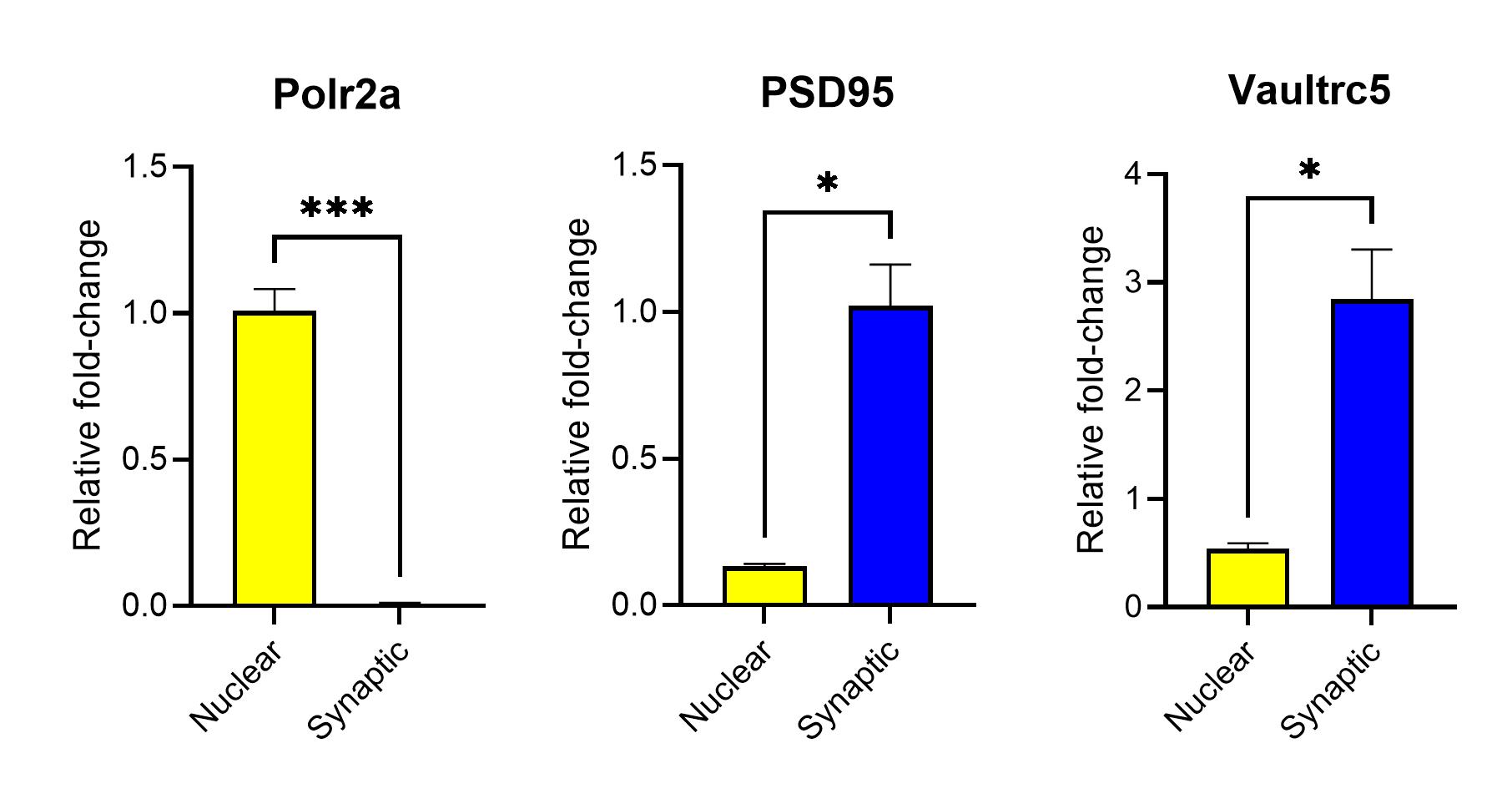

**A	B	C**
