## Supplementary Figure 4 for "Dynamic regulation of neuronal vault trafficking and RNA cargo by the noncoding RNA, Vaultrc5"

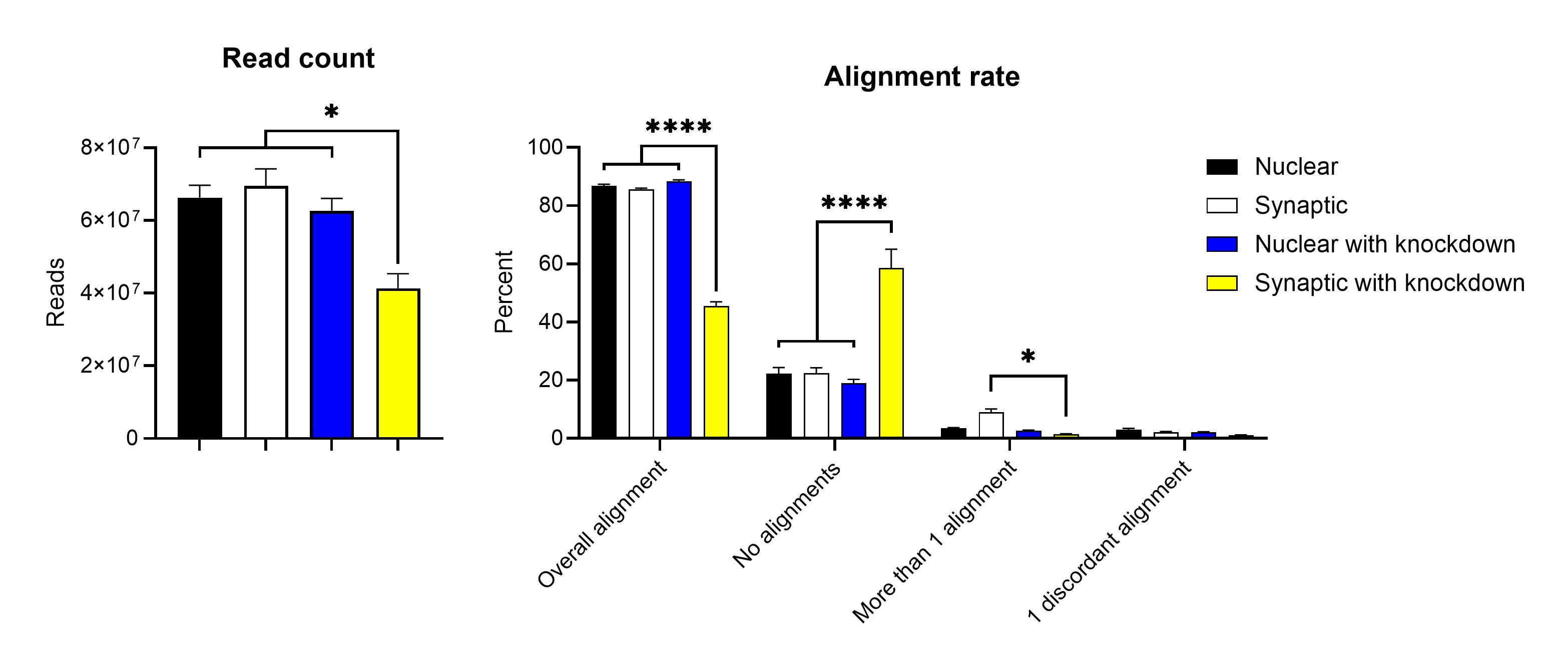

**Supplementary Figure X: HISAT2 quality control of sequencing libraries obtained from vault immunoprecipitation after Vaultrc5 knockdown.** A) Read count. Vaults sourced from the synaptic compartment after Vaultrc5 knockdown had significantly less reads than vaults sourced from the nuclear compartment (24903014 ± 5252344 more reads, p = 0.0157), synaptic compartment (28194961 ± 6110590 more reads, p = 0.0179), and nuclear compartment with knockdown (21298906 ± 5256672 more reads, p = 0.0324). B) Alignment rates. Synaptic with knockdown also had a significantly lower alignment rate than the nuclear (41.43 ± 2.645 % more), synaptic (40.16 ± 2.645 % more), and nuclear with knockdown (42.93 ± 2.645 % more) groups. Synaptic vaults had significantly more reads with multiple alignments than synaptic with knockdown (7.478 ± 2.645%). These data indicate that synaptic vaults with knockdown had fewer valid reads, and more unaligned reads than any other group, and that knockdown produced a minor, significant decrease in reads with multiple alignments in synaptic vaults. The tight clustering of individual samples throughout indicates that this is a group effect, supporting the idea that Vaultrc5 knockdown has a ‘gear shift’ effect on synaptic vaults.

**A	B**
